# Optogenetic control of receptors reveals distinct roles for actin- and Cdc42-dependent negative signals in chemotactic signal processing

**DOI:** 10.1101/2021.04.03.438340

**Authors:** George R. R. Bell, Esther Rincón, Emel Akdoğan, Sean R. Collins

**Affiliations:** Department of Microbiology and Molecular Genetics, University of California, Davis

## Abstract

During chemotaxis, neutrophils use cell surface G-Protein Coupled Receptors (GPCRs) to detect chemoattractant gradients^1–4^. The downstream signaling system is wired with multiple feedback loops that amplify weak inputs and promote spatial separation of cell front and rear activities^1, 5–8^. Positive feedback could promote rapid signal spreading^9^, yet information from the receptors is transmitted with high spatial fidelity, enabling detection of small differences in chemoattractant concentration across the cell^1^. How the signal transduction network achieves signal amplification while preserving spatial information remains unclear. The GTPase Cdc42 is a cell-front polarity coordinator that is predictive of cell turning, suggesting an important role in spatial processing^10^. To directly measure information flow from receptors to Cdc42, we paired zebrafish parapinopsina, an optogenetic GPCR that allows reversible ON/OFF receptor control with a spectrally compatible red/far red Cdc42 FRET biosensor. Using this new toolkit, we show that positive and negative signals downstream of G-proteins shape a rapid, dose-dependent Cdc42 response. Furthermore, F-actin and Cdc42 itself provide two distinct negative signals that limit the duration and spatial spread of Cdc42 activation, maintaining output signals local to the originating receptors.

## Results and Discussion

In leukocytes, chemotaxis is driven almost exclusively by GPCRs coupling to the Giα-family of G-proteins^1–4^. Through partially understood pathways, these receptors trigger activation of polarity and motility signaling driven by Rho-family GTPases, phospholipid signaling, and different actin assemblies. Rac, Cdc42, phosphatidylinositol (3,4,5)-trisphosphate (PIP3), and branched actin coordinate the cell front while RhoA and contractile actomyosin complexes define the cell rear^1, 5–8^. Accurate cell steering requires that the polarity programs receive and rapidly incorporate directional cues from receptors, enabling responses to differences in input strength across the cell. Prior studies indicate that receptors are uniformly distributed on the plasma membrane^11, 12^, and in *Dictyostelium discoideum* amoeba chemotaxis, G-protein activity largely mirrors receptor binding^4^, suggesting that spatial processing occurs downstream. In neutrophils, Cdc42 likely plays a key role in this process, as it stabilizes cell front/rear polarity, and asymmetry in its activity is predictive of cell turning^10, 13, 14^.

Signal processing in chemotaxis balances two potentially competing challenges. It must amplify signals using positive feedback to polarize cells with asymmetric protein activities, but it must also retain information about receptor status locally. Positive feedback could quickly distort spatial information, as it is capable of generating activity waves that can propagate faster than diffusion^9^. Indeed, Cdc42 activity can form traveling waves in neutrophil-like cells when actin is depolymerized^10^. Thus, many models for directional sensing in chemotaxis involve balancing of positive and negative feedback or feedforward loops that collaborate to restrict, but also amplify receptor-derived signals^2, 3, 5^. Nevertheless, the negative signaling mechanisms helping to maintain spatial information remain unclear.

We aimed to determine how inputs are processed downstream of receptors, including how signals spread spatially. Making these measurements requires sharply localized receptor activation, which would be very difficult to achieve with native attractants due to their rapid diffusion. Therefore, we developed parapinopsina, a nonvisual opsin GPCR, as an optogenetic tool that enables local (∼1 µm) and reversible stimulation of chemotaxis behavior through activation of Giα-family G-proteins. UV light activates parapinopsina by photo-isomerizing its 9-*cis*-retinal cofactor to *trans*-retinal, while green light (> 530 nm) inactivates the receptor and regenerates 9-*cis*-retinal^15, 16^, allowing rapid activation and deactivation cycles (Fig. 1a). Previously, OPN1SW (human blue cone opsin) and lamprey paparinopsina were shown to elicit a chemotactic-like response in mouse macrophage RAW 264.7 cells, supporting the use of optogenetic GPCRs to drive chemotaxis-like responses^17, 18^. In a sister article, we used this optogenetic tool to investigate overwriting cell front-rear polarity in neutrophils migrating in 1-D microfluidic channels, further demonstrating its usefulness for investigating complex signaling cascades^19^.

**Figure 1.**
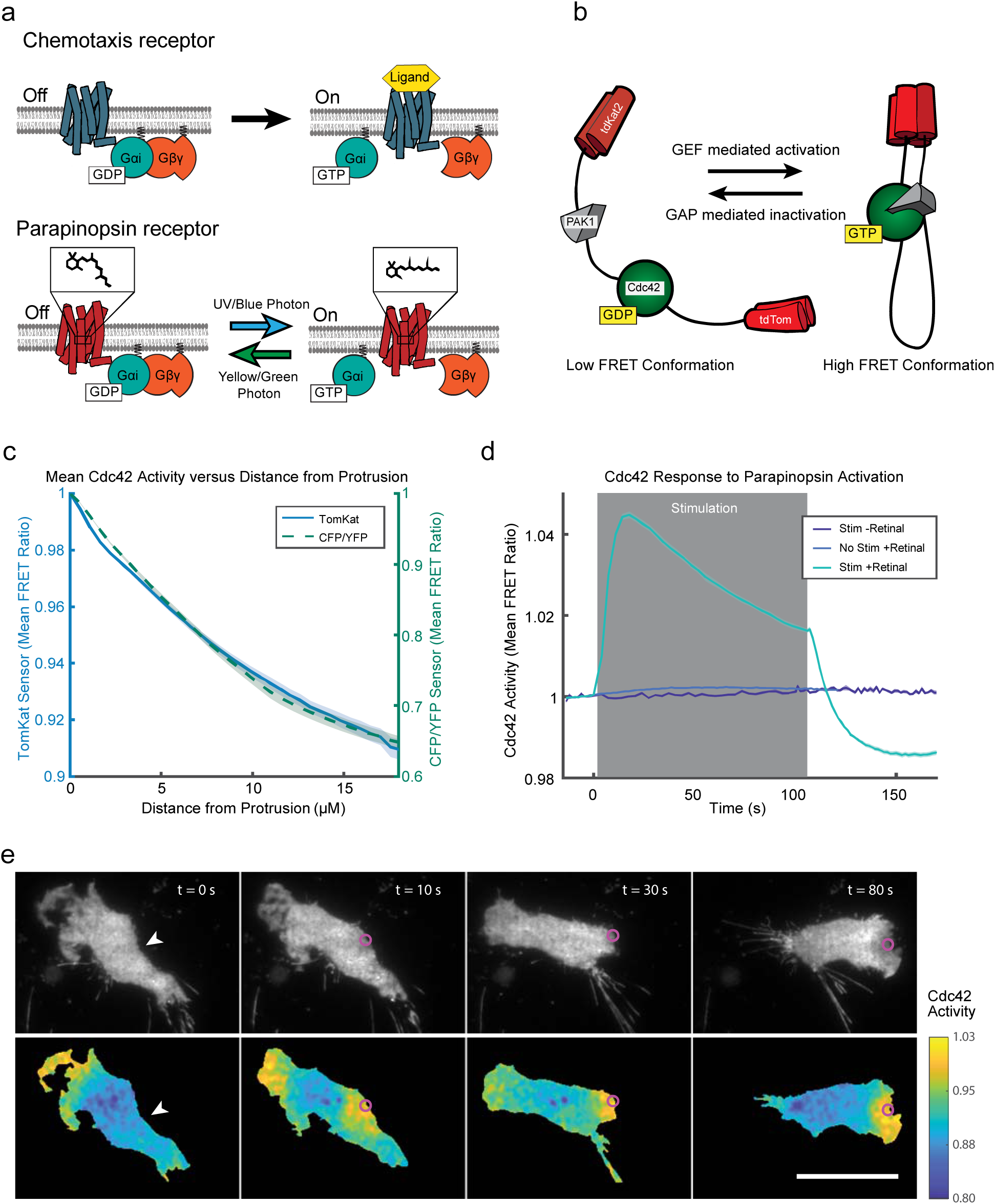
A molecular toolkit for optical control of receptor activity and measurement of signaling outputs in the same cell. (**a**) Like chemoattractant receptors (above), parapinopsina (below) is a Giα coupled GPCR that is activated by blue light. The chemical structure above the parapinopsina receptor represents the retinal chromophore that facilitates photon detection and receptor activation. Blue light photo-isomerizes 9-*cis*-retinal to all *trans*-retinal, activating the receptor. Longer wavelengths inactivate the receptor and photo-isomerize the retinal chromophore back to the cis-conformation. (**b**) Schematic for the “TomKat” FRET sensor that is spectrally compatible with the optogenetic GPCR (parapinopsina). The FRET donor is tdTomato while tdKatushka2 is the FRET acceptor. The sensor contains the Cdc42 binding domain from PAK1 and a C-terminally truncated Cdc42 that are separated by a linker domain. (**c**) The spatial activity profiles reported by TomKat and CFP/YFP FRET sensors in randomly moving differentiated PLB-985 cells. Shaded error regions represent ± s.e.m. of n=73 cells for TomKat sensor, n=59 cells for CFP/YFP sensor. (**d**) Cdc42 activity responses to global optogenetic GPCR activation are dependent on blue light stimulation and 9-*cis*-retinal cofactor. The response rapidly attenuates after stimulations cease, indicating that the receptor is inactivated by imaging with longer (> 530 nm) wavelengths of light. Shaded error regions represent ± s.e.m. of nwell replicates = 19 well replicates for no retinal condition (Stim-Ret), nwell replicates = 31 for no stimulation condition (No Stim +Ret), nwell replicates = 59 for stimulation with retinal (Stim +Ret). Relative light intesnsity = 10. Time on the x-axis is relative to the last FRET image before stimulation. (**e**) Focal stimulation of optogenetic-GPCR can repolarize a cell and drive a chemotaxis response. The white arrowheads indicate the target region pre-stimulus, while the magenta circles indicate the stimulated region. The Cdc42 TomKat sensor can be used to measure subcellular Cdc42 activity in the optogenetic GPCR stimulated cells. Scale bar, 25 µm.

To measure Cdc42 signaling downstream of parapinopsina in single cells, we modified an existing Cdc42 FRET biosensor^20^ to use a novel td-Tomato/td-Katushka2 (TomKat) FRET pair that is compatible with UV parapinopsina stimulation (Fig. 1b). Small GTPases are activated by guanine nucleotide exchange factors (GEFs) that catalyze GDP to GTP exchange, and inactivated by GTPase-activating proteins (GAPs) that promote GTP hydrolysis^21^. Importantly, this sensor reports on the local balance of GEF and GAP activity regulating Cdc42 (Fig. 1b). We validated the TomKat FRET sensor by comparing its spatial activity pattern to that of the original CFP/YFP FRET sensor in randomly migrating neutrophil-like cells (differentiated PLB-985 cells). Both sensors reported very similar spatial activity (Fig. 1c). Although the dynamic range for the TomKat sensor (∼11%) is less than that of the CFP/YFP sensor (∼56%), the TomKat sensor brightness still enables collection of high signal-to-noise data.

We next tested whether we could measure changes in Cdc42 activity downstream of parapinopsina, and sought to verify wavelength-dependent, reversible control of the receptor. In a population of cells, we monitored Cdc42 activity and applied a global, 100 second stimulation period in which we delivered pulses of UV light immediately after acquiring each FRET image. Stimulation triggered a rapid Cdc42 response that peaked in less than 20 seconds and remained elevated while UV light was applied. Once the UV stimulation ceased, the response attenuated immediately, consistent with long wavelength illumination inactivating receptors (Fig. 1d and Supplementary Video 1). Importantly, the response was dependent on UV-light stimulation, and exogenous 9-*cis* retinal.

We also verified that the receptor directed cell migration by stimulating a small (∼1-4 µm) region of the cell edge using a ∼2 µW spot of 407 nm light while recording Cdc42 activity. Local activation triggered cell repolarization and migration in the direction of stimulation (Fig. 1e and Supplementary Video 2). In optogenetically driven cells, Cdc42 activity was high at the leading edge, and decreased towards the cell rear, consistent with the spatial pattern observed in chemotaxing PLB-985 cells^10^. Collectively, these results demonstrate that we can optically activate parapinopsina to drive directed cell migration while recording spatial and temporal activity of Cdc42 using the spectrally compatible TomKat FRET sensor.

Many models for directional sensing and polarization involve integration of positive and negative signals downstream of receptors^3, 22, 23^. Therefore, we investigated whether both types of regulation act on Cdc42, by characterizing the temporal dynamics of receptor-initiated Cdc42 responses. (Fig. 2a). Using the global stimulation assay, we delivered a single stimulating light pulse to a population of cells, titrating the light stimulus strength. In all cases, the Cdc42 activity rapidly increased, peaked, and then attenuated, overshooting the pre-stimulus baseline within about 10 seconds (Fig. 2b and Supplementary Fig. 1a and Supplementary Video 3). The response eventually recovered from the negative overshoot, returning to a level near the pre-stimulus baseline after ∼2 min for the highest intensity stimulation (Supplementary Fig. 1b). Both the positive and negative phases of the response were dose dependent. Cdc42 is rapidly activated downstream of receptors, but this activation is countered by a slower, longer lasting negative regulation.

**Figure 2.**
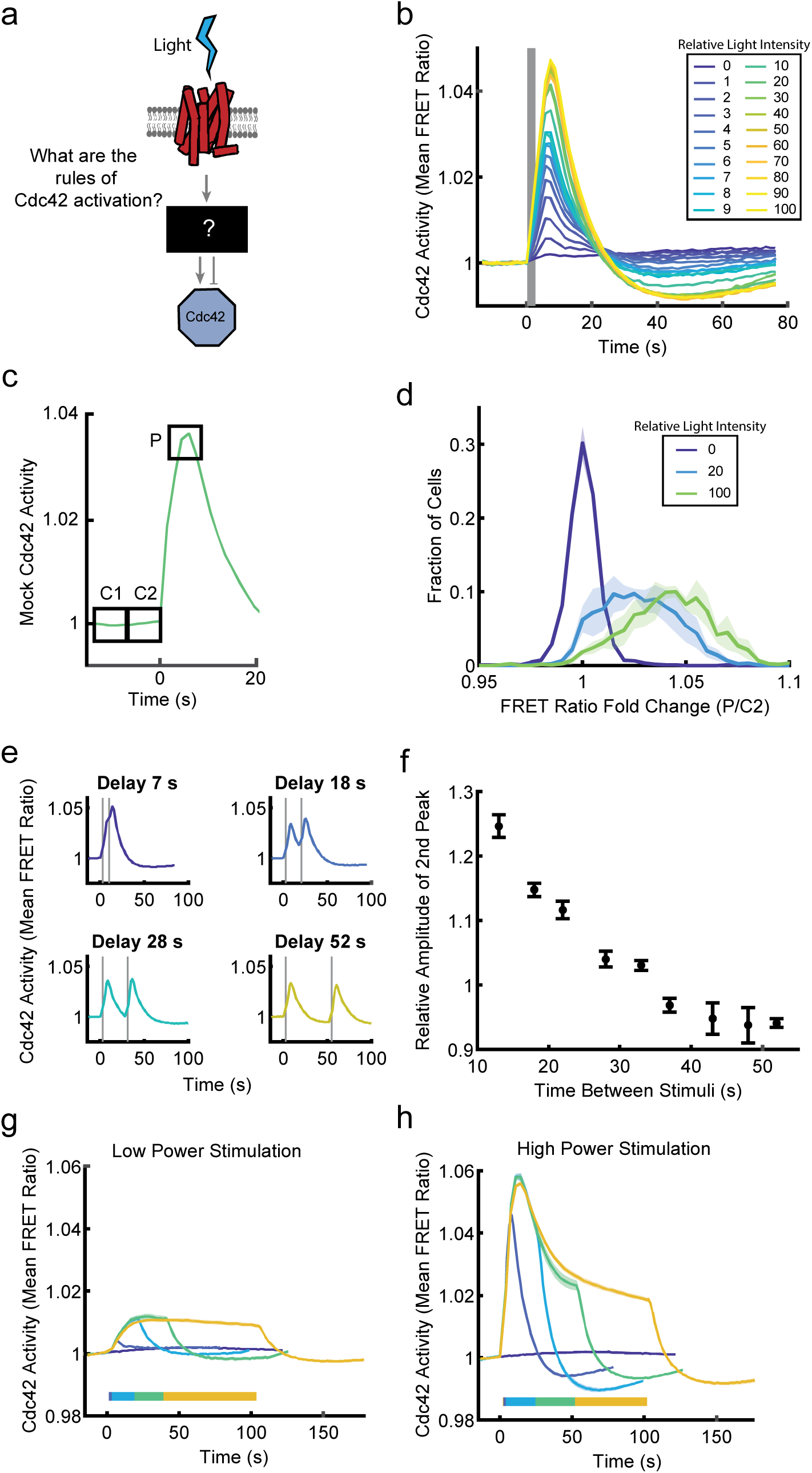
Dose-dependent positive and negative signals downstream of receptors shape a graded Cdc42 response. (**a**) Schematic of unknown signal processing between receptors and Cdc42. (**b**) Populations of PLB-985 cells expressing parapinopsina and the Cdc42 TomKat sensor were stimulated with a single light pulse of the indicated intensities. Mean FRET ratio was measured as a function of time. Stimulus duration is indicated by the gray bar. A minimum of seven well replicates were performed for each condition. See Supplementary Fig. 1a for plot with error bars. Tens to hundreds of cells were imaged in each well. (**c**) Schematic for analysis of single cell response amplitudes. For each cell, the response was broken into three, 7.5 s windows. The mean FRET ratio was calculated for each window (C1 = −13.5 s to −7.5 s, C2 = −6 s to 0 s, P = 4.5 s to 10.5 s). (**d**) Histograms of single-cell response amplitudes to a single light pulse stimulus of the indicated intensities. FRET ratio fold change was calculated by taking the ratio of the peak response (P) to the control window (C2). Total cell numbers of n=4127 cells (Relative light intensity = 0), n=2317 cells (Relative light intensity = 20), n=2261 cells (Relative light intensity =100) were analyzed from 4 independent experiments. The mean and standard error of the mean of the histogram values over the 4 experiments are shown. (**e**) Cdc42 responses are shown four different two pulse stimulation protocols with the indicated delay times between stimulations. The Relative Light Intensity was 10 for all plots. A minimum of four technical well replicates were performed for each condition. See Supplementary Fig. 2 for all two pulse stimulation plots. Stimulus duration indicated by gray bars. (**f**) Relative amplitude of the second Cdc42 response peak as a function of time between stimuli. The relative amplitude is calculated as the ratio of the second peak to the first peak. Time delay of 7 seconds was not included because a second peak could not be resolved from the first. The error bars indicate ± s.e.m. of nwell replicates = 6 for 13 s, nwell replicates = 6 for 18 s, nwell replicates = 6 for 22 s, nwell replicates = 6 for 28 s, nwell replicates = 26 for 33 s, nwell replicates = 4 for 37 s, nwell replicates = 4 for 43 s, nwell replicates = 4 for 48 s, nwell replicates = 4 for 52 s. (**g-h**) Repeated global stimulations were applied to simulate continuous receptor stimulation. Stimulation duration indicated by horizontal color bars. (**g**) Cdc42 response to the indicated stimulus durations with low power stimulation (Relative Light Intensity = 1). Shaded error regions represent ± s.e.m. of nwell replicates = 65 for non-stimulated, nwell replicates = 21 for 1 stimulation, nwell replicates = 20 for 7 stimulations, nwell replicates = 21 for 15 stimulations, and nwell replicates = 27 for 30 stimulations. (**h**) Cdc42 response to the indicated stimulus durations with high power stimulation (Relative light lntensity = 100). Shaded error regions represent ± s.e.m. of nwell replicates = 65 for 0 stimulation condition, nwell replicates = 12 for 1 stimulation, nwell replicates = 12 for 7 stimulations, nwell replicates = 12 for 15 stimulations, and nwell replicates = 77 for 30 stimulations.

The fact that the Cdc42 response was tunable (Fig. 2b) was intriguing since the chemotaxis signaling network is known to contain positive feedback loops that amplify responses, promote polarity, and contribute to “excitable” system behaviors, which can include all-or-nothing responses, refractory periods, and propagating waves of activity^7, 24, 25^. Cdc42 is regulated by positive feedback through PAK1^26^ and its activity generates traveling waves when the actin cytoskeleton is depolymerized^10^, indicating some excitable systems features. To verify that the Cdc42 response is tunable, despite positive feedbacks, we analyzed the same experiments at the single cell level, where cell-to-cell variability could not obscure all-or-nothing behavior. For each cell, we measured the response amplitude, relative to baseline, at the typical peak response time (Fig. 2c,d); as a control we performed the same analysis using a pre-stimulus time window (Supplementary Fig. 1c). We found that for intermediate stimulus levels, most cells responded with intermediate response amplitudes, indicating that the response was titratable at the single-cell level (Fig. 2d). While this contrasts with earlier observations of switch-like behavior for PIP3 responses in HL-60 cells^27^, the different roles of Cdc42 and PIP3 at the cell front may require alternate regulation mechanisms. Additionally, our system provides very precise control of stimulus intensity and timing, resolving differences in responses that peak within 10 seconds.

Leveraging precise control of cell stimulation, we used two-pulse and prolonged stimulation protocols to understand how the cells integrate signals temporally, potentially revealing adaptive behavior or other features of negative regulation. By applying two, pulse stimulations with varying time delays, we found that responses to sequential stimuli were largely independent. Closely spaced inputs added to produce larger responses, with no obvious refractory behavior (Fig. 2e,f and Supplementary Fig. 2). Interestingly, this result differs from Ras activation in Latrunculin-A treated *Dictyostelium* cells and PIP3 in HL-60 cells where a refractory period was observed^27, 28^. The lack of a refractory period suggests that the Cdc42 circuit can rapidly respond to new inputs, a feature likely important for responding dynamically to pathogen cues and navigating complex environments. Next, we applied prolonged stimulations to investigate potential adaptive behavior. In response to prolonged low-power stimulus, the Cdc42 response gradually increased (dependent on continued input), until it reached a plateau at about 20 seconds, and then only slightly attenuating until the stimulation was removed (Fig. 2g). In contrast, a stronger prolonged stimulus caused a response that rapidly peaked, and then quickly began to attenuate. However, the rapid attenuation did not cause adaptation, but gave way to a shoulder phase with slower attenuation until the stimulation ceased (Fig. 2h). These experiments reinforce that the Cdc42 response is graded based on receptor input strength, but they also suggest complex, multi-tiered negative regulation of the circuit with differential kinetics.

Ultimately, we wanted to connect features of receptor-mediated Cdc42 signaling dynamics with molecular components to understand signal processing. Since Cdc42 activity is regulated by multiple feedback connections^13, 26, 29^, we reasoned that this feedback may play an important role in shaping Cdc42 dynamics. To test this, we generated a clonal, homozygous Cdc42 knockout (Cdc42-KO) cell line using CRISPR/Cas9 ribonucleoprotein complexes targeted to excise a ∼100 bp region of exon 4. We validated the knockout using amplicon sequencing (Supplementary Fig. 3a) and western blot (Fig. 3a and Supplementary Fig. 3b). To examine the loss of feedback at the cellular level, we assessed Cdc42-KO cells for migration and polarity defects (Fig. 3b-g). Qualitatively, randomly migrating Cdc42-KO cells tended to make multiple cell fronts that were less stable than the controls cells (Fig. 3e,f and Supplementary Video 4). Furthermore, we observed reduced migratory persistence in randomly migrating cells, as evident in downward curvature of a mean square displacement plot (Fig. 3b), and in the faster decay of directionality of migrating cells (Fig. 3c). The multiple cell front behavior and poor migratory persistence are consistent with observations from HL-60 cells expressing dominant-negative Cdc42^13^ and Cdc42-null mouse neutrophils^14^. Additionally, we noticed that many Cdc42-KO cells formed cell fronts that pulled away from the cell body, stretching out thin cytoplasmic tethers (Fig. 3g and Supplementary Video 5). Quantifying this behavior, we found that ∼40% of Cdc42-KO cells formed cytoplasmic tethers compared to about 4% of control cells (Fig. 3d). Tethers observed in control cells were also typically much shorter and thicker than in the Cdc42-KO line. Interestingly, inhibition of myosin-II with blebbistatin can cause cytoplasmic tethers HL-60 cells^3^, and local Cdc42 activation can induce a long-distance myosin response^29^. Collectively, these findings suggest that Cdc42 cell front activity may mediate long-range regulation of protrusion-inhibiting, cell rear polarity signals.

**Figure 3.**
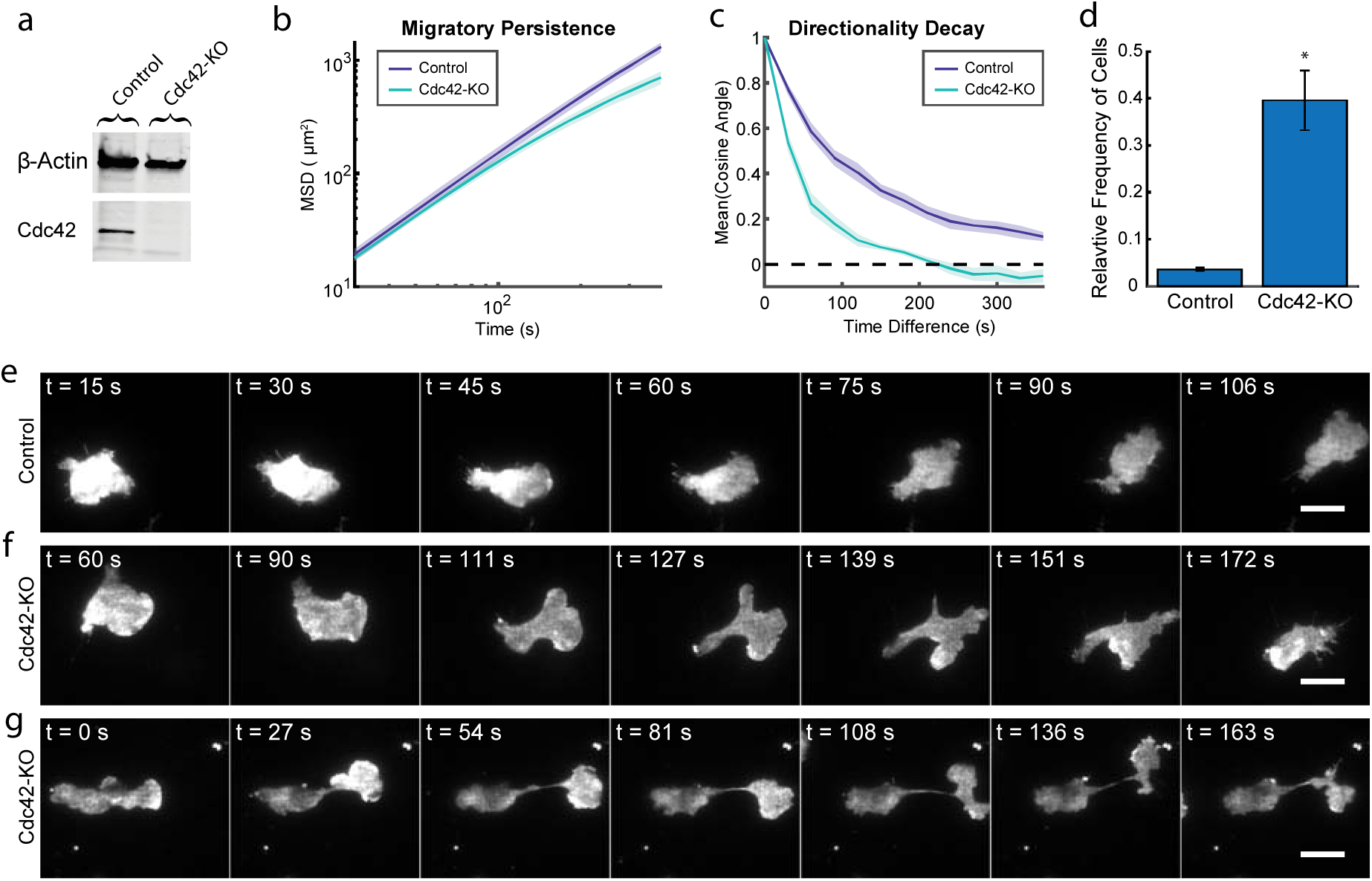
Genetic knockout of Cdc42 reveals a central role in organizing and maintaining cell polarity. (**a**) Western blot comparing the Cdc42 knockout (Cdc42-KO) and control cell lines stained for β-actin and Cdc42. (**b**) The mean squared displacement (MSD) was measured as a function of time for control and Cdc42-KO cells randomly migrating in an under-agarose condition. Shaded error regions represent ± s.e.m. of nexperiment = 4. A total of 10,687 control cells and 3,018 Cdc42-KO cells were analyzed across the four experiments. (**c**) As a measure of directional persistence, the mean cosine of the angle between a migrating cell’s movement direction at two different time points was measured as a function of the difference in time between the two measurements. Cells migrating in a straight line would have a mean cosine angle of 1. Shaded error regions represent ± s.e.m. of nexperiment = 4. A total of 4,821 control cells and 1,098 Cdc42-KO cells were analyzed across the four experiments. (**d**) Fraction of cells exhibiting the cytoplasmic tether phenotype. The error bars represent ± s.e.m. of nexperiment = 4. A total of 331 control cells and 194 Cdc42-KO cells were analyzed across the four experiments. * indicates p-value =0.0108 using t-test with unequal variances. (**e**) Time-lapse image series of control cell randomly migrating. Control cells tend to maintain one cell front at a time and migrate more persistently. Time points match the corresponding supplemental video. Scale bar, 15 µm. (**f**) Time-lapse image series of a Cdc42-KO cell randomly migrating. This cell displays a range of phenotypes including: a very large, crescent-shaped leading edge in the first time point and multiple repolarization events thereafter. Time points match the corresponding supplemental video. Scale bar, 15 µm. (**g**) Time-lapse image series of a representative Cdc42-KO cell randomly migrating. This cell displays the cytoplasmic tether phenotype where the leading edge pulls away from the cell body, but remains linked by a thin cytoplasmic filament. Time points match the corresponding supplemental video. Scale bar, 15 µm.

Next, we probed the components and signaling processes that shape the Cdc42 response using drug perturbations and the Cdc42-KO cell line, with particular interest in the complex negative regulation. First, we investigated whether negative regulation involves heterotrimeric G-protein independent mechanisms, as documented in *Dictyostelium* amoeba^27^ (Fig. 4a). We used pertussis toxin (PTX) to inhibit Giα family G-protein signaling and asked whether the negative phase of the Cdc42 response was left intact. Instead, we found that PTX dramatically suppressed both the positive and negative phases, indicating that both are dependent on Giα family G-proteins (Fig. 4b,c).

**Figure 4.**
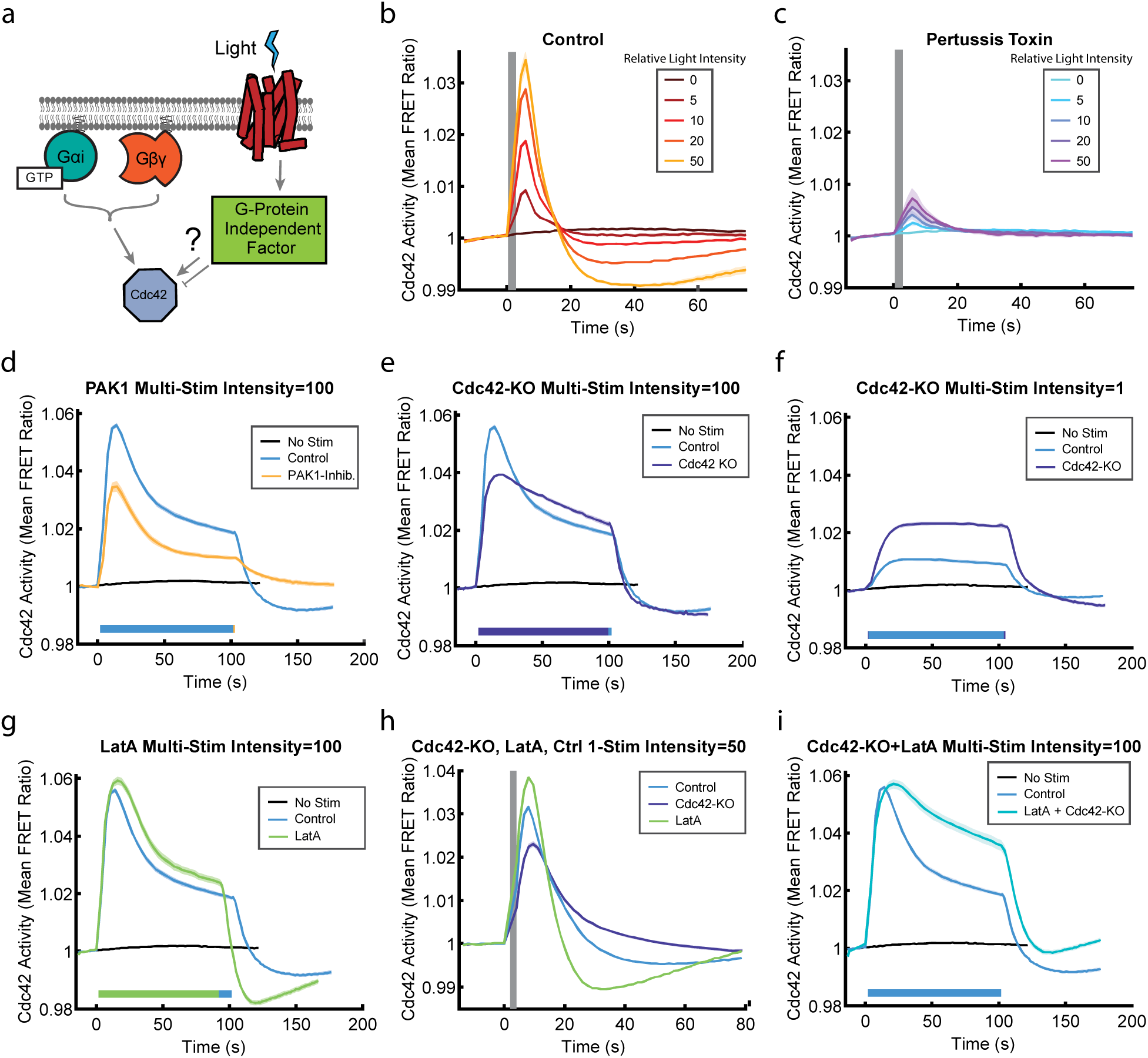
Multiple negative signals downstream of G-proteins collaborate to regulate the Cdc42 response. (**a**) Schematic indicating the potential for Cdc42 response regulation independent of heterotrimeric G-protein signals. (**b-c**) Single pulse stimulations of populations of PLB-985 cells expressing parapinopsina and the Cdc42 TomKat sensor either without (b) or after treatment with 600 ng/ml Pertussis toxin (PTX) to inhibit Giα dependent signals (c). Shaded error region represents ± s.e.m of nwell replicates = 16 for all conditions. Stimulus duration indicated by gray bar. (**d**) Comparison of Cdc42 responses to prolonged stimulation with or without treatment with 5 μM PAK1 inhibitor IPA-3. Shaded error region ± s.e.m. of nwell replicates = 65 for non-stimulated, nwell replicates = 77 for control, nwell replicates = 32 for PAK1 inhibited. Relative light intensity =100. Stimulation duration indicated by color bars. (**e**) Comparison of Cdc42 TomKat sensor response measurements under the same conditions for Cdc42-KO cells and control cells. Shaded error region ± s.e.m. of nwell replicates 65 for non-stimulated, nwell replicates = 77 for control, nwell replicates = 56 for Cdc42-KO. Relative light intensity =100. Stimulation duration indicated by color bars. (**f**) Comparison of the responses of the same cell lines to lower power stimulations. Shaded error region ± s.e.m. of nwell replicates = 65 for non-stimulated, nwell replicates = 27 for control, nwell replicates = 16 for Cdc42-KO. Relative light intensity =1. Stimulation duration indicated by color bars. (**g**) Comparison of Cdc42 responses to prolonged stimulation with or without treatment with 1 μM of the actin depolymerizing agent Latrunculin A. Shaded error region represents ± s.e.m. of nwell replicates = 65 for non-stimulated, nwell replicates = 77 for control, nwell replicates = 32 for Latrunculin-A-treated cells. Relative light intensity =100. Stimulation duration indicated by color bars. (**h**) Comparison of single-pulse stimulation responses for the same conditions as in (g). Shaded error region ± s.e.m. of nwell replicates = 69 for control, nwell replicates = 42 for Cdc42-KO, nwell replicates = 64 for Latrunculin-A-treated cells. Relative light intensity = 50. Stimulus duration indicated by gray bar. (**i**) Comparison of responses for untreated control cells and Cdc42-KO cells treated with 1 μM Latrunculin-A. Shaded error region represents ± s.e.m. of nwell replicates = 65 for non-stimulated, nwell replicates = 77 for control, nwell replicates = 32 for Latrunculin-A + Cdc42-KO cells. Relative light intensity =100. Stimulation duration indicated by color bars.

Next, we sought to verify that we could detect changes in the Cdc42 response due to specific perturbations downstream of the receptor and G-proteins. Since PAK1 kinase amplifies Cdc42 signaling through a positive feedback loop,^26^ we reasoned that inhibition of PAK1 would result in a lower Cdc42 response magnitude. As expected, PAK1 inhibition with IPA-3 reduced the Cdc42 response overall (Fig. 4d and Supplementary Fig. 4a). Importantly, the kinetic profile shape was largely the same, indicating that we can detect alterations in the Cdc42 response that are due to specific disruptions in the signaling pathway.

We then investigated Cdc42-dependent feedback using the Cdc42-KO background. We hypothesized that if a Cdc42-dependent feedback was the primary pathway controlling signal attenuation, then it should be disrupted by loss of endogenous Cdc42. Since the Cdc42 TomKat sensor monitors the balance of regulating GEF and GAP activities, we expected its signal to remain elevated after stimulation. Instead, we observed more complex dynamics indicating multiple regulatory pathways. With a strong stimulus, the Cdc42-KO response magnitude was reduced as with PAK1 inhibition (Fig. 4d), suggesting that the Cdc42-dependent positive feedback loop was impaired. However, the attenuation dynamics were also slower and lacked the initial fast phase (Fig. 4e). Further supporting an important role for negative feedback, the Cdc42-KO response magnitude was double that of the control in response to a weaker stimulus (Fig. 4f). Interestingly, the post-stimulation attenuation phase of the response was unaltered by the knockout. Collectively, these results indicate that Cdc42 activity is negatively regulated by Cdc42-dependent feedback and by Cdc42-independent mechanisms.

We reasoned that actin assembly could be a second negative regulator, as it is known to feed back to polarity signaling through membrane and cortical tension. Growing protrusions increase tension, which globally limits actin polymerization and Rac activity^30, 31^. To test this, we treated cells with Latrunculin-A to depolymerize F-actin. We found that the Cdc42 response dynamics retained fast and slow attenuation phases in response to prolonged stimulation. However, the amplitude was increased, the initial attenuation phase was slightly slower, and the post-stimulation negative response was faster and stronger (Fig. 4g). These differences were more obvious in pulse stimulation experiments (Fig. 4h). These results suggest that rapid, actin-dependent signaling limits Cdc42 responses, but that other major negative signals are independent of F-actin and membrane tension. Finally, we treated Cdc42-KO cells with Latrunctulin-A to determine if both were functioning in the same pathway. The response combined features of both perturbations, resulting in slower signal attenuation than with either alone (Fig 4i). These results suggest that F-actin and Cdc42-dependent feedback mechanisms provide two distinct negative signals regulating Cdc42.

These two negative regulators emerged as likely candidates for spatially constraining the spread of signaling downstream of receptor inputs. To test this, we developed a total internal reflection fluorescence (TIRF) microscopy assay to directly measure spatial signal processing in the basal plasma membrane. We automated cell identification and delivery of either one or five, low powered micron-scale, stimulus pulses to the cell center and followed the response over time (Fig. 5a and Supplementary Video 6). Qualitatively, the response was rapid, but did not spread across the whole cell (Fig. 5a,b). We quantified this by measuring the mean change in FRET ratio as a function of distance from the stimulation site (Fig. 5c). To simplify our analysis, we analyzed only non-polarized and slow-moving cells. In control cells, the Cdc42 response to a single local stimulus rapidly peaked (3.8 s) and began attenuating, returning to the pre-stimulus baseline at ∼10 s (Fig. 5d and Supplementary Fig. 5a). Notably, the response did not spread across the whole cell, remaining nearly constant in regions distal (> 6 µm) to the stimulation site. However, the results were different for Cdc42-KO and Latrunculin-A-treated cells. Immediately after stimulation (0.8 s), all three conditions were indistinguishable, indicating that the profile of receptor activation was the same (Fig. 5e). In contrast, Cdc42-KO and Latrunculin-A-treated cells had larger, prolonged responses that extended more than 8 µm from the stimulation site at the response peak (Fig. 5f, Supplementary Fig. 5b-d). Because the single pulse stimulation experiment is transient, we asked whether the Cdc42 circuit could restrict information spread in the context of prolonged (5-pulse) stimulation. While the control Cdc42 response was still locally restricted near the stimulation site (Fig. 5g), the signal spreading in the Latrunculin-A condition was enhanced, highlighting the requirement for F-actin for proper spatial signal processing (Fig. 5h). Collectively, these results suggest that the Cdc42 circuit is organized to spatially restrict the spread of information from the receptor, while rapid response attenuation limits the duration of the signaling event.

**Figure 5.**
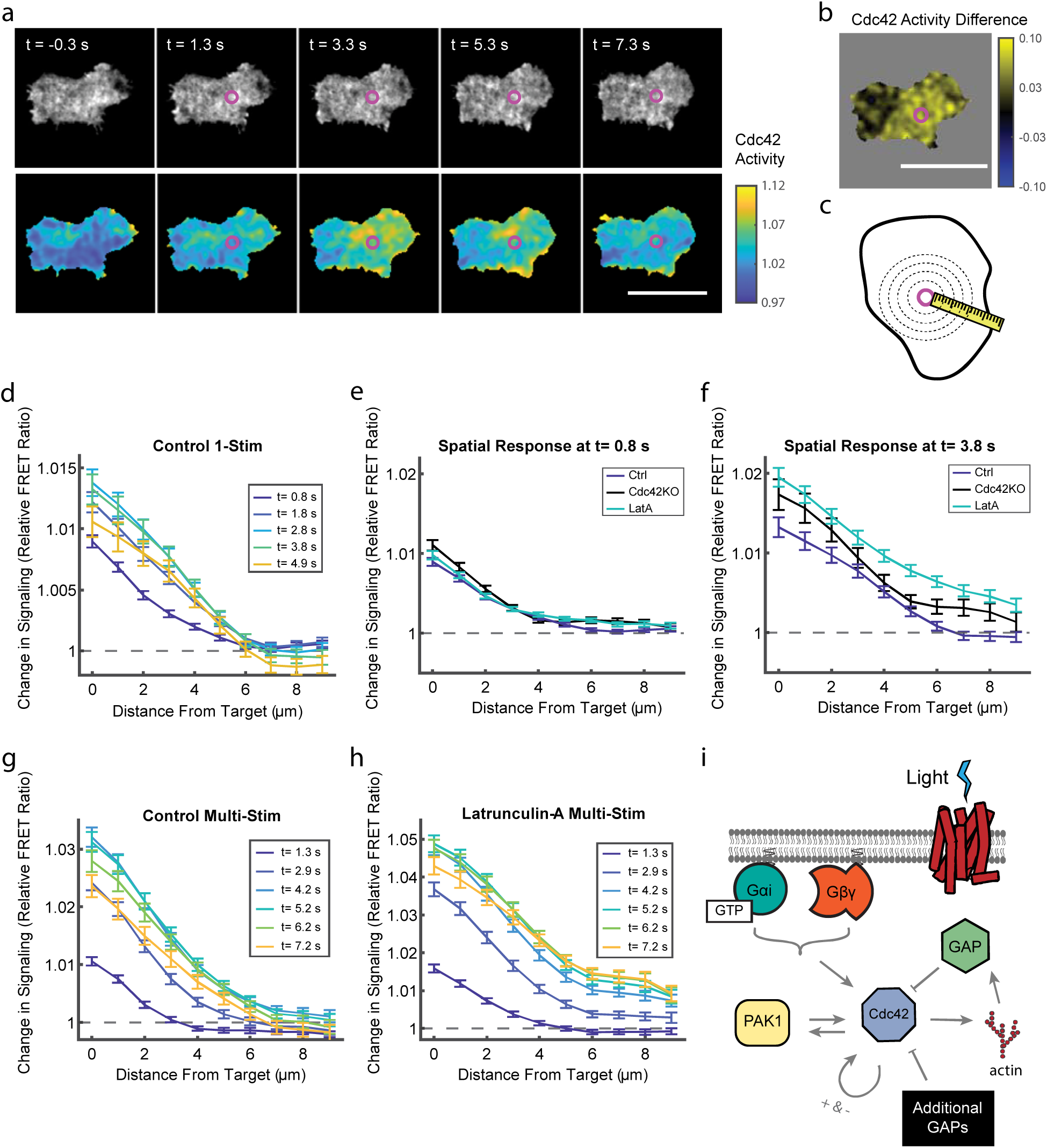
F-actin and Cdc42 spatially constrain the spread of signals downstream of receptors. (**a**) A control cell responding to the center stimulation experiment. A single laser pulse (4.3 µW, 10 ms duration) was applied between frame 1 and 2. Purple circle indicates the stimulation site. Top panel images are the sum of the two FRET channels. Bottom panel images are FRET ratio images. Times relative to stimulation are indicated. Scale bar, 15 µm. (**b**) The spatial Cdc42 response was calculated as the difference between the FRET ratio images before and after stimulation (Frame 3 – Frame 1) for the cell shown in (a). Scale bar, 15 µm. (**c**) Schematic for center stimulation experiment analysis strategy. Cell pixels were aggregated based on their distance from the stimulus target site (magenta circle) for each frame in the experiment. (**d**) Relative Cdc42 response as a function of distance from the stimulus target site for control cells at the indicated time points. One 0.8 µW light-pulse of 10 ms duration was delivered at t = 0s. Error bars represent ± s.e.m. of n=181 cells. (**e-f**) Relative Cdc42 response as a function of distance from stimulus target site for control, Cdc42-KO, and Latrunculin-A conditions at t=0.8 s (**e**) and t=3.8 s (**f**) post-stimulation. Error bars represent ± s.e.m. of n=181 cells for control, n=67 for Cdc42-KO, n=175 for Latrunculin-A-treated cells. (**g-h**) Relative Cdc42 responses as a function of distance from the stimulation site for control and Latrunculin-A conditions in response to five sequential 0.8 µW light-pulses of 10 ms duration delivered immediately after successive images, with the first stimulus delivered at t = 0s. Error bars represent ± s.e.m. of n=142 cells for control, n=102 for Latrunculin-A-treated cells. (**i**) Schematic indicating the positive and negative regulators of the Cdc42 response identified in this study.

How spatial information encoded by the chemotaxis receptors is processed by the signal transduction network has remained a longstanding open question. Using a new optogenetic molecular toolkit that enables precise measurements, we show that the Cdc42 signaling circuit is optimized to limit the duration and spatial spread of responses downstream of the receptor. Loss of either F-actin or Cdc42-dependent feedback loops were sufficient to disrupt the response’s short and local, spatiotemporal span. In particular, the rapid negative regulation from F-actin was surprising, as F-actin also participates in a positive feedback loop with Rac and PIP3^32–34^. The negative regulation could depend on a number of GAPs that interact with F-actin^35^. GAPs often have complex regulation that require multiple signaling inputs; thus, actin binding could play a role in GAP activation or positioning^21, 35^. More generally, our results demonstrate that GAP activity, in addition to GEF activity, is actively regulated downstream of receptor activation, and that multiple negative signals cooperate to coordinate the Cdc42 response (Fig. 5i). Of particular interest are the unidentified GAPs and regulatory mechanisms that respond quickly after receptor activation to attenuate and restrict responses. Finally, the Cdc42-KO phenotypes for both maintenance of cell-wide polarization and local preservation of information downstream of receptor inputs contributes to a growing body of evidence that Cdc42 plays a central role in integrating directional inputs with the cell polarity cascade.

## Acknowledgements

The authors would like to thank Amalia Hadjitheodorou for insightful discussions of image analysis methodologies as well as edits to the manuscript. Additionally, we thank members of the Collins lab for support, especially Dean Natwick, Kwabena Badu-Nkansah, and Sam Hayes for thoughtful manuscript edits and discussions. This work was supported by an NIH Director’s New Innovator Award (DP2HD094656) to S.R.C. G.R.R.B would like to thank the National Science Foundation Graduate Research Fellowship (Grant Number 1650042) for support.

## Author Contributions

G.R.R.B and S.R.C conceived of the project. G.R.R.B constructed the parapinopsina and TomKat control cell lines and optimized the parapinopsina system. E.R.G. generated the Cdc42-KO clonal cell line, while E.A. created Cdc42-KO lines expressing parapinopsina and the TomKat FRET sensor. G.R.R.B conducted the experiments and performed the data analysis with guidance from S.R.C. G.R.R.B and S.R.C. interpreted the results and wrote the manuscript. All authors have read and edited the manuscript.

## Methods

### Reagents

This study used Latrunculin-A at a final concentration of 1 µM (Calbiochem Cat# 428021), the PAK1 kinase inhibitor, IPA-3 (5 µM final concentration, Cayman Chemical Cat# 14759), and pertussis toxin (600 ng/ml final concentration, Invitrogen Cat# PHZ1174). Latrunculin-A and IPA-3 were reconstituted in Dimethyl Sulfoxide (DMSO), while the pertussis toxin was diluted in sterile, distilled water.

### Cloning

A human codon-optimized version of the zebrafish parapinopsina gene was printed using the Thermofisher Gene Art service in a pUC57 bacterial expression plasmid. The Gene Art product also contained the prolactin signal sequence peptide on the N-terminus of the parapinopsina gene to enhance protein expression on the cell membrane^36, 37^. Using Gibson cloning, the prolactin-parapinopsin construct was C-terminally tagged with mCitrine and inserted into a lentiviral vector.

The Cdc42 TomKat FRET sensor was created by modifying the previously described CFP/YFP Cdc42 FRET sensor^20^. Using a combination of traditional restriction and Gibson cloning, the fluorescent proteins from the original sensor were removed and replaced with td-Tomato as the FRET donor and td-Katushka2 as the FRET acceptor. td-Katushka2 was a gift from Michael Davidson (Addgene # 56049)^38^. All plasmids and plasmid maps will be made available on Addgene for the published version of this manuscript.

### Cell culture

PLB-985 cells were cultured in RPMI 1640 (Gibco) complete media as previously described^39^. Cells were differentiated into a neutrophil-like state by culturing 2 x 10^5^ cells per ml in RPMI 1640 with 4.5-5% FBS, 100 mg/ml streptomycin/ 100 U/ml penicillin (P/S), 1.3% DMSO, and 2% Nutridoma-CS (Roche) for 6 days^39^. HEK-293T (ATCC CRL-11268) were used for lentiviral production. Cells were cultured in high glucose DMEM (Sigma-Aldrich, D5671) that was supplemented with 10% FBS, 1% Glutamax and 100 mg/ml streptomycin/ 100 U/ml penicillin (P/S). All cell lines were maintained in an incubator at 37 °C and 5% CO2. For imaging experiments, a modified “L-15 imaging media” (Leibovitz’s L-15 media lacking dye, riboflavin, and folic acid (UC Davis Biological Media Services) was used to minimize media autofluorescence.

### Cell line construction

The Cdc42-TomKat FRET sensor plasmid contains the Inverted Terminal Repeats (ITR) of the piggybac transposon system^40^. To create a stable cell line, the Cdc42-TomKat FRET sensor plasmid was co-electroporated at a 1:1 ratio with the piggybac transposase expression plasmid. Electroporation was achieved with the Neon electroporation system (Sigma-Aldrich). 2 x 10^6^ cells were resuspended in the R-buffer from the kit, then 5 µg of each plasmid were added to the cells. Cells were electroporated in the 100 µL pipettes provided in the kit at 1350 V for 35 ms.

The parapinopsina construct was then stably integrated into PLB-985 cells expressing the Cdc42-TomKat sensor using 2^nd^ generation lentivirus. To produce the virus, HEK-293T cells were co-transfected with envelope, packaging and transfer plasmids using Mirus TransIT-2020 (Cat# MIR 5404) transfection reagent. The following ratio of plasmids was used to transfect 1 well on a P6 plate: 0.64 µg Envelope: 1.26 µg Packaging: 1.93 µg Transfer.

Cdc42-KO cells were created by electroporating CRISPR-Cas9 Ribonucleoprotein (RNP) complexes into the PLB-985 cell line. A pair of CRISPR guides targeting a 102 bp region of Cdc42’s exon 4 were designed and purchased from Synthego as part of their Gene Knockout Kit v2. Guide 1 had a sequence of 5’-TTTCTTTTTTCTAGGGCAAG while Guide 2 was 5’-ATTTGAAAACGTGAAAGAAA. Purified Cas9 protein was purchased from the QB3 MacroLab at UC Berkeley. To generate the RNPs, a solution containing 180 pmoles of the Synthego guide mixture and 5 µM Cas9 diluted in the Neon R-buffer was incubated at 37 °C for 10 min then stored at RT. Electroporation was conducted using a suspended-drop electroporation device as described^41^. For this device, a maximum volume of 10 µL is electroporated per well on a P96-well plate. Cells (3e^5^) were resuspended in 5 µL of Mirus Ingenio electroporation buffer (MIR 50111), mixed with 5 µL of RNP solution, and then electroporated at 120 V for 9 ms. Cells were allowed to recover in RPMI complete media with 20% FBS for 1 week. To check heterogeneous knock out efficiency, genomic DNA (gDNA) was harvested, and PCR was used to amplify a fragment that covered 300 bp upstream and downstream of the CRISPR guides. This PCR fragment was sanger sequenced and then analyzed using Synthego’s Inference of CRISPR Edits (ICE) tool. The Cdc42 forward sequencing primer had an estimated indel frequency of 83% while the reverse sequencing primer indel frequency was 93%. Based on these results, we proceeded to clonal analysis. Serially diluted cells were plated at a density of 0.5 cells/well on a P96-well plate. Wells containing a single cell were identified by phase-contrast microscopy and tracked as the culture expanded. Ten clones were selected and were evaluated using several methods. First, the clones were analyzed using the Synthego ICE tool. Promising clones were next assessed by Amplicon sequencing, and Quantitative Western blot.

### Amplicon sequencing

gDNA was harvested from 5 million cells for WT and Cdc42-KOclones (Invitrogen Purelink Genomic DNA Kit). The purified gDNA was then used as a PCR template for primers that flank the CRISPR cut site. The forward primer sequence was 5’-ACACTCTTTCCCTACACGACGCTCTTCCGATCTccagcatgcttttaacactttgagg while the reverse primer sequence was 5’-GACTGGAGTTCAGACGTGTGCTCTTCCGATCTgaaaggagtctttggacagtggtg. Upper case letters in the primer indicate the partial Illumina adapter sequences. The PCR product was cleaned (Zymo DNA clean and concentrator-5) and then sent to Genewiz for 2 x 250 bp amplicon sequencing (Amplicon-EZ service). The amplicon sequences were analyzed using MATLAB to identify unique sequences after excluding sequences that did not match the primer sequences. Additionally, a small number of nonspecific sequences of less than 100 bp that were observed in all samples were removed. For each remaining unique sequence, the number of identically matching reads was counted. Sequences were aligned to the genomic sequence of the human Cdc42 gene to determine deleted or mutated regions. The 3 most frequently observed sequences for both forward and reverse sequences are shown (Supplementary Fig. 3a).

### Immunoblots

Cdc42 protein expression levels were compared using Western Blotting. Differentiated PLB-985 cells were lysed at 4 °C in NP40 buffer (150 mM NaCl, 50 mM Tris Base, 1% NP40, PH 8.0) plus protease inhibitor (Thermo cat#: 78429) by repeatedly passing the cells through a 21-gauge syringe needle. Cell lysates were centrifuged to remove cellular debris, snap frozen in liquid nitrogen, and stored at −80 °C. Lysates were thawed on ice, and protein levels were quantified with the Pierce 660 nm protein absorbance assay kit (22660) on a Molecular Devices SpectraMax spectrophotometer. Lysates were then solubilized by mixing 3:1 lysate to Li-Cor 4x loading buffer (928-40004) plus 10% Beta-Mercaptoethanol (Sigma). Samples were denatured by boiling for 5 min followed by cooling on ice for 2 min. 15 µg of total protein per sample were loaded on a 4-15% gradient polyacrylamide gel (BioRad #4561084) and run for 15 min at 100 V followed by ∼35 min at 150 V. Proteins were transferred onto nitrocellulose membranes for 1.25 hrs at 100 V. The membrane was blocked using Li-Cor Intercept TBS blocking buffer for 30 min. Anti-Cdc42 primary antibody (Abcam Cat# ab187643) was diluted 1:10,000 in blocking buffer and incubated overnight at 4 °C on a rotary shaker. The following day, the membrane was washed 3X with TBST (1% Tween 20 in Tris Buffered Saline), and then incubated with secondary antibody at RT for 1 hr on a rotary shaker in the dark. The secondary antibody (Li-Cor IRDye® 800CW Donkey anti-Rabbit IgG Secondary Antibody) was diluted 1:15,000 into blocking buffer + 0.2% Tween 20. Post staining, the membrane was washed 3x in TBST. Membranes were washed once in deionized water and imaged using the Li-Cor Odyssey Imager (model 9120) using the 800 nm channel. Post imaging, the antibody labeling steps were repeated on the same membrane for the rabbit anti-β-Actin antibody (Cell Signaling Technology Cat# 4967). The β-actin antibody was diluted 1:1000 in blocking buffer, incubated overnight, and labeled the next day with the Li-Cor 800CW Donkey anti-rabbit secondary. The membrane was imaged using the Li-Cor Odyssey Imager at 800 nm (Supplementary Fig. 3b).

### Retinal preparation

All retinal solutions were prepared in a dark room with red-light sources. 9-*cis*-retinal (Sigma Aldrich R5754) was dissolved in argon purged, 200 proof ethanol (Sigma Aldrich) to reach a concentration of 10 mg/mL. Aliquots were stored in small amber glass tubes (Sigma Aldrich) at −80 °C. 9-*cis*-retinal is hydrophobic, and requires a 1% w/v BSA carrier solution^42, 43^. To prepare the carrier solution, Bovine Serum Albumin (BSA), Fraction V—Low-Endotoxin Grade (Gemini Bio 700-102P) was dissolved in L-15 imaging media. Importantly, this BSA product is low in non-esterified fatty acids, which can inhibit the effectiveness of the BSA as a carrier^43^. Next, 10 µL of retinal stock was diluted to a working concentration of 10 µg/ml by incrementally adding the 1% BSA solution (9 x 10 µL, 4 x 100 µL, 1 x 500 µL, 9 x 1 mL) until the final volume was 10 mL. The working retinal solution was then stored in a light-proof box and mixed overnight at 4 °C. Prior to an imaging experiment, cells were resuspended in the 10 µg/mL retinal solution and incubated for 1hr at 37 °C. The diluted retinal solution was kept for up to 3 days. All downstream processing steps after cells were incubated in retinal were carried out in the dark.

### Microscope configuration for TomKat FRET sensor imaging

All imaging experiments were conducted using a Nikon Eclipse Ti stand with dual Andor Zyla 4.2 sCMOS cameras. The microscope is controlled by MATLAB via Micromanager, allowing automated and highly repeatable experimental scripts. Simultaneous image acquisition for each FRET channel was achieved using a Cairn TwinCam LS image splitter. The TomKat FRET sensor requires 561nm excitation of td-Tomato while the FRET acceptor (td-Katushka2) has its excitation maxima at ∼590nm. Thus, we designed our optics to isolate the td-Tomato emission spectra (combined ∼573-600 nm bandpass) to one camera, while the td-Katusha2 and any potential td-Tomato bleed-through emissions were sent to the second camera (> 610 nm). For global stimulation experiments with the TomKat FRET sensor, populations of cells were imaged at 20x (Nikon Apochromat 0.75 NA) via epifluorescent illumination with the X-Cite XLED1 GYX LED. For the high spatial resolution under-agarose experiments, a 561 nm laser line (Andor) was used for Total Internal Reflection Fluorescence Imaging (TIRF) with a 60x Nikon Apochromat objective (1.49 NA). Our GYX LED has a dual 560/640 band excitation filter, thus the TomKat filter cube requires a short pass excitation filter that blocks the longer wavelengths (Semrock, BSP01-633R excitation filter). The TomKat filter cube uses a single edge dichroic that reflects ∼560 nm light while passing longer wavelengths for emission (ZT561rdc Chroma). Finally, the cube uses two ∼570 nm long pass emission filters to prevent the TIRF laser illumination from reaching the cameras (ET570lp Chroma). The TwinCam image splitter uses the following filter configuration to properly capture the two FRET channels on their appropriate cameras. The image splitter dichroic is positioned to reflect wavelengths < 605 nm to one camera and pass longer wavelengths to the other (ZT594rdc Chroma). The td-Tomato side of the cube (< 605 nm) additionally uses an emission filter to further block wavelengths > 600nm to ensure that only td-Tomato emission signal is hitting the camera (ET560_80m Chroma). Finally, the td-Katushka2 side of the cube uses a long pass emission filter that blocks wavelengths < 610 nm.

### Camera and illumination corrections

The dark-state noise for each camera was empirically measured by capturing 79 images without illumination and with the light path switched to the oculars. The median over the stack was used to generate the dark-state correction image, which is then subtracted from all experimental images. Next, uniform dye preparations were used to correct for variability in pixel responsiveness as well as camera and illumination artifacts. Rose bengal dye solution (0.3 mg/mL) was centrifuged at 21,000 RCF for 5 min to remove insoluble particles. 2 µL of dye was plated in the center of a P96-glassbottomed imaging plate (Cellvis P96-1.5H-N). A 5 mm round coverslip was applied to the dye droplet to create a thin, uniform layer of dye. TomKat epifluorescent dye images were captured (n ≥ 1000) and the median for each camera was taken over the image stack. To correct for differences between each half of the camera sensor, the ratio of the mean pixel intensity for two rows above and below the middle of the camera chip was used to generate a correction factor. This correction factor was then applied to the bottom half of an image-sized matrix of ones, creating a half-chip correction. The half-chip correction is multiplied by all imported images after the dark-state subtraction. Dust in the microscope light path was observed to cause a small dark spot in the dye image on one of the cameras. To correct for this illumination artifact, the median dye image was smoothed using a broad gaussian filter (sigma = 30) to create a filtered image that does not have the spot artifact. The ratio of the median dye image to the smoothed dye image was used to generate a logical mask for the dim pixels in the artifact spot. The pixels in the mask were smoothed with a gaussian filter (sigma = 10), then this image was divided by the median dye image to create a dust correction image. These images were generated for both cameras and were applied after the half-chip correction for images collected at 60x.

A gradient in FRET ratio activity was empirically observed from the top to bottom of the TomKat FRET sensor images. A ratio correction image was developed to remove this gradient in FRET activity for both the 20x and 60x objectives. Images of unstimulated Cdc42 TomKat FRET sensor cells were collected with cells positioned randomly throughout the images so that at least one cell was imaged on every portion of the camera sensor. These images were processed with our standard pipeline to generate FRET ratio images. The pixel-by-pixel median FRET ratio was then taken over all images (including only data from pixels inside cells). To reduce noise and local variability, this median image was then broken into 24×24 pixel blocks and the median was taken for each block. The resulting image was smoothed using a gaussian filter (sigma = 5) and the image was resized to match the input image size. To apply the correction, FRET ratio images are divided by the ratio correction image.

For center stimulation experiments, local activation of the parapinopsina receptor with 407 nm light focused through the FRAP module causes a small amount of photobleaching of the Cdc42 FRET sensor which could cause bias in signaling measurements (Supplementary Fig. 6). To correct for this, we measured the diameter and recovery rate for bleached sensor molecules using a high-power stimulus (37 µW for 10 ms) to develop a diffusion model of the bleached sensor using an empirically fit initial bleaching pattern. The diffusion coefficient used in the model was 0.5 µm/s. A bleaching correction was generated for each post-stimulation frame for each cell by modeling sensor diffusion, centered around the empirically measured FRAP stimulation target site and scaled to match the experimental laser stimulation power.

### Global stimulation imaging and image analysis

Post retinal incubation, differentiated cells were resuspended in L-15 imaging media that contained 2% FBS and were plated in glass-bottomed 384-well plates at a density of ∼1100 cells/µL x 20 µL (Corning Cat#: 4581). All imaging experiments were conducted at 37 °C and were terminated after 5 hours of imaging. Separate cell lines or treatment conditions were positioned so that all lines were imaged evenly throughout the experiment. Global UV stimulations were delivered using the DAPI epifluorescent channel. Stimulation power was manipulated by altering exposure time and LED power. For continuous stimulation experiments, UV stimulation and TomKat FRET sensor imaging were alternated until the stimulation period was over. Images were captured at a frame rate of 1.5 s, although prolonged global stimulation experiments required DAPI and TomKat filter cube switching, which slowed the image capture rate.

For the global stimulation experiments, pertussis toxin (600 ng/ml), IPA-3 (5 µM) and Latrunculin-A (1 µM) were used to perturb signaling pathways. The pertussis toxin was added to the retinal incubation step, and the incubation duration was extended to 2.5 hrs. The control cells received the same extended retinal duration. For the Latrunculin-A and IPA-3 conditions, the inhibitors were added to the retinal solution and incubated for 1 hr. Additionally, cells were plated with these compounds added to the imaging media to ensure inhibition during the experiment.

FRET pair images were aligned with a custom MATLAB function that uses a coordinate-mapping strategy as described^10^. Aligned images were cropped to ensure that both images are the same size. Next, the dimmest pixels (1.5 percentile) across all frames were used to define the background pixels. Since the cells are densely packed, empty-well images (median of ∼1400 for each channel) were used to estimate the background spatial profile. The dimmest pixels were used to scale the brightness of the empty-well images prior to background subtraction. Next, pixels were filtered and removed from further analysis if they were dim, near saturating, or if the FRET ratio of the pixel was low (< 0.8), indicating that the cell was unhealthy or dead. Dying and dead cells have high autofluorescence that is independent of the FRET sensor, causing high FRET donor signal and low FRET values. Because this filter was applied on a per pixel basis that changed across time points, a conservative dead cell FRET ratio threshold was selected. Finally, the mean intensity was calculated for each channel before the ratio was taken. Whole-frame mean FRET ratio values were computed for each well at each timepoint. Plots were generated from means and standard errors computed over all replicate wells from all experiments. Qualitative trends were consistent across experiments from different days. For global stimulation plots, time on the x-axis is relative to the Cdc42 FRET image immediately preceding stimulation.

To investigate the Cdc42 response on a single cell level, single-pulse, global stimulation experiments were re-examined. Images were aligned, cropped, and background subtracted as above. The sum of the FRET donor and acceptor images was used for masking as the sum has better signal to noise ratio and is less susceptible to changes in signal intensity due to FRET. To generate cell masks for tracking, the sum image was smoothed using a gaussian filter with a standard deviation of 2 and then sharpened using unsharp masking. Automatic thresholding was used to define the cell masks. Strict minimum and maximum cell area thresholds were applied to the masks to remove cell fragments or cell aggregates. Finally, the centroids of the cell masks were tracked across all frames using a reciprocal nearest neighbor tracking strategy. Cells that could not be tracked across all images were removed from further analysis. Dead cells were filtered on a per cell basis using the FRET ratio mean of the first ten frames (pre-stimulus). Based on a bimodal distribution of measured baseline FRET values, cells with a baseline FRET ratio < 0.95 were thus removed from the analysis. Additionally, cells that contained saturated pixels in either the td-Tomato or td-Katushka2 channel were removed. FRET ratio fold change was assessed by breaking the cellular response into three, 7.5 s windows (Control1 = −13.5 s to - 7.5 s, Control2 = −6 s to 0 s, Peak = 4.5 s to 10.5 s). The response mean was taken over the five frames in each window and then the ratio of the Peak window to the Control2 window was taken. Additionally, the ratio of Control1 to Control2 was computed (Supplementary Fig. 1c). Histograms for single-cell data were computed for 4 experiments conducted on different days. These histograms were averaged, and the standard error of the mean was computed for each histogram bin. The qualitative trend was the same for all four experiments.

### Under-agarose cell preparation

Several experiments were conducted using an under-agarose cell preparation on a 96-well plate format (Cellvis Cat#: P96-1.5H-N). A detailed protocol for the under-agarose preparation can be found as described^44^. In brief, differentiated cells (∼1,000 – 1,500) were plated in the center of a well in a 5 µL drop of 2% FBS + L-15 imaging media. Cells were allowed to adhere to the glass for 5 min, before a 195 µL layer of 1.5% low-melt agarose (Invitrogen Cat#: 16520-100) mixed with 10% FBS + L-15 imaging media equilibrated to 37 °C was overlayed on top. The agarose solution was allowed to solidify at RT for 40 min before the plate was transferred to the microscope incubator and warmed to 37 °C for 40 min prior to imaging. Importantly, the cells must remain in the interface between the agarose and the glass for proper motility. Thus, careful pipetting of the agarose solution is required to prevent dislodging cells from the glass.

### TIRF image background subtraction and cell segmentation

Raw images were first corrected for the camera dark-state noise, differences in camera chip sensitivity, and dust in the light path as described above. FRET pair images were aligned using the coordinate-mapping strategy noted above. Next, cells were segmented by first summing the FRET donor and acceptor images to enhance signal to noise. The sum images were used to conservatively define background and cell object pixels. Next, background intensity images were computed using the median intensity of background pixels in the local neighborhood for each pixel. Background images were subtracted from the sum image, and object edges were enhanced using unsharp-masking. To perform unsharp-masking, the image was smoothed using a broad gaussian filter (sigma=25), and then was subtracted from the original image. Finally, the cell object masks were defined using Otsu’s threshold method.

After segmentation, each FRET donor and acceptor image was background subtracted using the background mask defined in the segmentation section. Next each image was smoothed using a gaussian filter (sigma=1) and pixels not in the cell mask were defined as not a number (NaN) to remove them from further analysis. The FRET ratio image was calculated as FRET acceptor divided by FRET donor. The FRET ratio image was then divided by the ratio correction image to account for the observed gradient in FRET sensor activity in the images.

### Cdc42 spatial activity analysis

The spatial activity pattern of the Cdc42 TomKat and CFP/YFP FRET sensors were measured and compared to validate proper function of the new TomKat FRET sensor. Differentiated Cells were plated using the under-agarose preparation. Then time-lapse TIRF microscopy images for randomly migrating, unstimulated cells were collected for each FRET pair. Assessment of the FRET sensor spatial activity was determined using the following image analysis steps as described^10^. Cells were segmented and tracked using the approximate nearest neighbor search method based on cell centroid positions. Cell tracks were manually curated to select only frames where cells were consistently moving. Cell protrusions were defined from frame to frame by subtracting the cell masks and identifying the largest connected protruding edge region. The protrusion was also required to be within one pixel of the defined protrusion from the previous and following frames. Next the shortest distance between each pixel in the cell mask and the protrusion mask was calculated using the bwdistgeodesic MATLAB function. The mean FRET ratio was calculated as a function of distance from the leading edge of the cell.

### Cdc42-KO cell phenotype characterization

Differentiated PLB-985 WT and Cdc42-KO cells were plated using the under-agarose preparation. Unstimulated, randomly migrating cells were imaged using TIRF microscopy. Cells were segmented as described above and grayscale movies were generated. Cells were manually counted as the cytoplasmic tether phenotype was difficult to accurately segment. Four experiments were analyzed, and the standard error of the mean was calculated for the mean of the four experiments.

### Analysis of persistence of cell migration

Differentiated PLB WT and Cdc42-KO cells were plated using the under-agarose preparation. Unstimulated, randomly migrating cells were imaged using 10X magnification epifluorescence microscopy. Cells were imaged for 7 min, with images acquired every 30 seconds. Cells were segmented and tracked using custom MATLAB software as previously described^45^. Briefly, cells were segmented using a manually determined intensity threshold, with minimum and maximum cell area thresholds. Cells were tracked using a reciprocal nearest neighbor algorithm. Two measures of persistence were computed. First, we computed the cosine of the angle between the direction of movement in the first 30 s, and the direction of movement in each subsequent frame-to-frame step. Only cells that moved at least 5 µm in the first 30 second step were included for analysis to capture only moving cells for which an initial direction could be determined accurately. We then computed the mean cosine value for each time point to determine the decay of directional persistence. Second, we computed the mean squared displacement as a function of time. Only cells that moved at least 5 µm from their starting position during the imaging interval were included for analysis to exclude unhealthy or nonmoving cells. The mean and standard error of the mean were computed at each timepoint over four independent experiments.

### Optogenetic laser stimulation using the FRAP module

Optogenetic stimulation was achieved for high resolution, TIRF microscopy experiments, by stimulating cells with a 407 nm laser (Coherent Cube) focused through a Fluorescence Recovery After Photobleaching (FRAP) module on the microscope. The TomKat dichroic can pass 407 nm light, allowing for rapid FRAP stimulation without changing the filter cubes. To focus the FRAP module, PLB cells expressing the TomKat FRET sensor were plated using the under-agarose preparation. Cells were imaged in TIRF to determine the appropriate focal plane. The FRAP laser was tuned to 40 ms exposure and 10 mW power. Cells were then selected and imaged using the FRAP channel. The X and Y translation knobs were used to adjust the FRAP spot until it was near the center of the image (∼512×512 on a 1024×1024 pixel image). Next the Z-adjustment knob was used to focus the FRAP spot into a tight gaussian. Power measurements of the FRAP laser at the objective were taken using a Thorlabs handheld optical power meter (PM100D) and microscope slide power sensor (S170C). Cell driving experiments were conducted using ∼2 µW power measured at the objective. Center-stimulation experiments were conducted using 4.3 or 0.8 µW power. The input laser could not be reliably tuned to achieve the 0.8 µW power value. Instead, a 1% neutral density filter was added to the FRAP module light path. This power level was also below the detection limit for the power meter. Three higher power measurements were collected with the ND filter installed, and a line was fitted to determine the nominal laser power required to stimulate cells at ∼0.8 µW. On each experimental day, ten pictures of the FRAP spot were collected and averaged to identify the pixel with the maximum FRAP spot intensity. This pixel location was saved and used for the remainder of the experiment. The mean FRAP spot image was saved and used for identifying the FRAP spot location in the image analysis steps.

### Optogenetic cell driving and cell center-stimulation assays

For optogenetic experiments, differentiated cells were pre-incubated with the 9-*cis-*retinal solution, and all plating steps of the under-agarose preparation were conducted in the dark with a red headlamp for illumination. For Latrunculin-A experiments, cells were treated with 1 µM Latrunculin-A during the 1 hr retinal incubation period. Additionally, Latrunculin-A was added to the 10% FBS + L-15 imaging media prior to mixing with the agarose solution to yield a final concentration of 1 µm Latrunculin-A.

Automated imaging scripts for cell driving and center-stimulation assays were developed in MATLAB. To automatically identify cells, an individual well on a 96-well plate was broken into a 7×7 search grid. A scan image was collected for each position in the grid, and objects were identified using the cell masking strategy described above. If a cell was detected, the script would center the stage on the cell using the cell centroid. Driving a cell required consistent delivery of FRAP stimulation to a region on the cell edge. To achieve this, a target angle was selected by the user. A MATLAB function was developed that would calculate the angle between the cell centroid and all outermost pixels on the cell perimeter. The perimeter pixel that had the closest match to the target angle was selected. The stage was translated so that the cell edge was centered on the empirically measured FRAP spot location and a low powered, ∼2 µw 407 nm FRAP stimulus was applied to the cell. Next the FRET images were collected, and the new cell coordinates were calculated from the td-Katushka-2 image using the cell masking and edge selection strategy. The stage coordinates for each movement required for imaging the cell were recorded and used for post processing of figures and movies. Cells were imaged using 3 or 5 second intervals. Once the experiment was completed for the individual cell, the script moved to the next position in the scan grid. When an individual grid was completed the script then moved to the next well and repeated the process. Depending on the experimental conditions, several hundred cells could be imaged in one, four-hour experiment automatically.

The center-stimulation assay used the same scanning and cell detection methods as the optogenetic driving assay. Once a cell was detected, the stage was moved to the centroid of the cell mask. This centering process was used for every frame before the FRAP stimulus was applied. Post stimulation however, the centering step was no longer required; this was true for both the single stimulation and 5-pulse stimulation experiments. Cells were imaged at a frame rate of 0.5 seconds. Prior to each center stimulation experiment, 3-5 cells were driven with the cell driving assay to confirm that the optogenetic receptor and stimulation system were functional.

To evaluate the spatial spread of Cdc42 activity in cells stimulated with the center-stimulation protocol, cell images were pre-screened to remove cell clusters or dead cells. Additionally, the saved FRAP spot image was converted into a logical mask where the brightest pixel in the FRAP spot was set to one. FRET images were aligned and segmented, and FRET ratio images were calculated using the TIRF image segmentation method described above. Next, cell mask, FRAP mask, and FRET images were adjusted using the stage translation recordings so that all image frames were aligned with the first frame in the set. For accurate analysis results, cells were screened using an automated custom MATLAB function to ensure limited translocation away from the FRAP stimulus site. The function removed cells if the FRAP mask could not be overlayed with the cell mask, if the FRAP mask was less than four pixels from the cell edge or if the Euclidean distance traveled for the cell centroid was greater than 4 µm for any frame post-stimulation. The photobleaching correction was applied to the FRET ratio images to account for FRET sensor bleaching due to FRAP stimulation. All pixels within a cell mask were measured to determine their distance away from the FRAP stimulation site pixel. These distance masks were used to sort pixels into concentric bins that were 1 µm in width. Pixels closest to the stimulation site were placed in bin 1 while pixels on the cell periphery were placed in the higher bins (Fig. 5c). The mean intensity values for the donor and acceptor were calculated for each bin, and then the FRET ratio for each bin was calculated. For this analysis, the ratio correction image was applied to the FRET acceptor images before the bin analysis was applied. These ratio bin measurements were compiled for all cells within experimental groups and used to generate the plots in Fig. 5.

### Statistical analysis

All error bar and shaded error regions represent standard error of the mean. Significance values were calculated using student’s t-test with unequal variance (MATLAB’s ttest2 function).

## Notes

### Competing Interest Statement

The authors have declared no competing interest.

