## Supplementary Information for "Optogenetic control of receptors reveals distinct roles for actin- and Cdc42-dependent negative signals in chemotactic signal processing"

Supplementary Figure 1

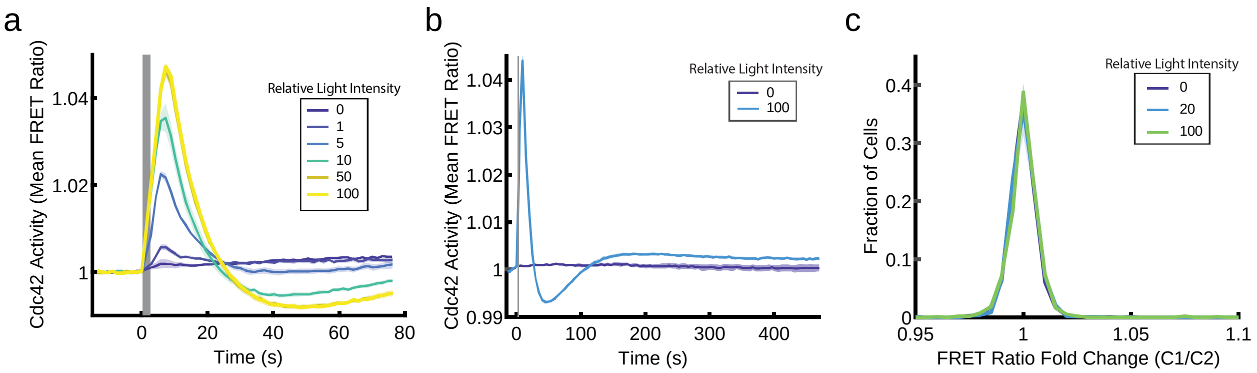


**Supplementary Figure 1:** Dose-dependent positive and negative signals downstream of receptors shape a graded Cdc42 response. (**a**) Subset of Cdc42-dose response curves, highlighting the graded nature of the Cdc42 response. Shaded error regions represent ± s.e.m. of $n_{\text{well replicates}}$ = 7 for each stimulation condition. (**b**) Single stimulation experiment with extended duration captures late phase oscillatory behavior as the Cdc42 response adapts to a baseline level. Shaded error regions represent ± s.e.m. of $n_{\text{well replicates}}$ = 11 for each stimulation condition. (**c**) Control for fraction of cells responding to light stimulus strength. Control for FRET ratio fold change was calculated by taking the ratio of the control window (C1) to the control window (C2). Shaded error regions represent ± s.e.m. of $n_{\text{experiment}}$ = 4. Across the four experiments, n=4127 cells (relative light intensity = 0), n=2317 cells (relative light intensity = 20), n=2261 cells (relative light intensity =100).

Supplementary Figure 2

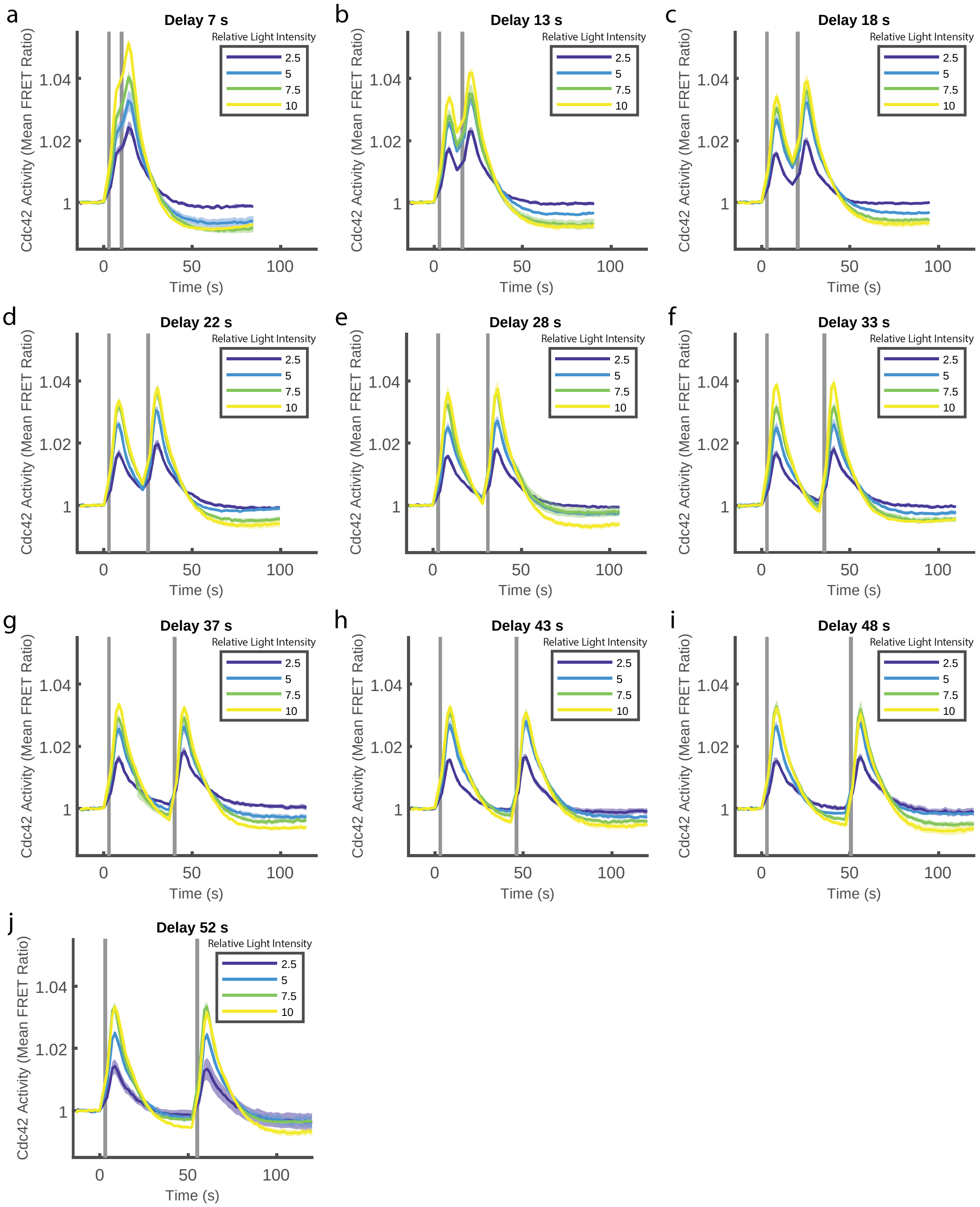

**Supplementary Figure 2:** Responses to sequential stimuli are independent. (**a-j**) Cdc42 response to two equal duration and light intensity stimulations with varying time delays. (**a**) Shaded error regions represent ± s.e.m. of $n_{well\ replicates}$ = 6 (Relative light intensity = 2.5), $n_{well\ replicates}$ = 6 (Relative light intensity = 5), $n_{well\ replicates}$ = 6 (Relative light intensity = 7.5), $n_{well\ replicates}$ = 30 (Relative light intensity = 10). (**b**) Shaded error regions represent ± s.e.m. of $n_{well\ replicates}$ = 7 (Relative light intensity = 2.5), $n_{well\ replicates}$ = 6 (Relative light intensity = 5), $n_{well\ replicates}$ = 6 (Relative light intensity = 7.5), $n_{well\ replicates}$ = 6 (Relative light intensity = 10). (**c**) Shaded error regions represent ± s.e.m. of $n_{well\ replicates}$ = 30 (Relative light intensity = 2.5), $n_{well\ replicates}$ = 7 (Relative light intensity = 5), $n_{well\ replicates}$ = 6 (Relative light intensity = 7.5), $n_{well\ replicates}$ = 6 (Relative light intensity = 10). (**d**) Shaded error regions represent ± s.e.m. of $n_{well\ replicates}$ = 6 (Relative light intensity = 2.5), $n_{well\ replicates}$ = 30 (Relative light intensity = 5), $n_{well\ replicates}$ = 7 (Relative light intensity =7.5), $n_{well\ replicates}$ = 6 (Relative light intensity = 10). (**e**) Shaded error regions represent ± s.e.m. of $n_{well\ replicates}$ = 6 (Relative light intensity = 2.5), $n_{well\ replicates}$ = 6 (Relative light intensity = 5), $n_{well\ replicates}$ = 30 (Relative light intensity = 7.5), $n_{well\ replicates}$ = 6 (Relative light intensity = 10). (**f**) Shaded error regions represent ± s.e.m. of $n_{well\ replicates}$ = 5 (Relative light intensity = 2.5), $n_{well\ replicates}$ = 5 (Relative light intensity = 5), $n_{well\ replicates}$ = 5 (Relative light intensity =7.5), $n_{well\ replicates}$ = 26 (Relative light intensity = 10). (**g**) Shaded error regions represent ± s.e.m. of $n_{well\ replicates}$ = 5 (Relative light intensity = 2.5), $n_{well\ replicates}$ = 4(Relative light intensity = 5), $n_{well\ replicates}$ = 4 (Relative light intensity = 7.5), $n_{well\ replicates}$ = 4 (Relative light intensity = 10). (**h**) Shaded error regions represent ± s.e.m. of $n_{well\ replicates}$ = 26 (Relative light intensity = 2.5), $n_{well\ replicates}$ = 5 (Relative light intensity =5), $n_{well\ replicates}$ = 4 (Relative light intensity = 7.5), $n_{well\ replicates}$ = 4 (Relative light intensity = 10). (**i**) Shaded error regions represent ± s.e.m. of $n_{well\ replicates}$ = 4 (Relative light intensity = 2.5), $n_{well}$

$_{replicates}$ = 26 (Relative light intensity = 5), $n_{well\ replicates}$ = 5 (Relative light intensity = 7.5), $n_{well\ replicates}$ = 4 (Relative light intensity = 10). (**j**) Shaded error regions represent ± s.e.m. of $n_{well\ replicates}$ = 4 (Relative light intensity = 2.5), $n_{well\ replicates}$ = 4 (Relative light intensity = 5), $n_{well\ replicates}$ = 26 (Relative light intensity = 7.5), $n_{well\ replicates}$ = 4 (Relative light intensity = 10).

### Supplementary Figure 3

**a**

Forward sequencing results

| | Synthetic sgRNA #1 | | Synthetic sgRNA #2 | # of additional bp not matching ref sequence |
|---|---|---|---|---|
| Reference Sequence | ATTTTTCTTTTTTCTAGGGCAAGAGG | ... | ATTTGAAAACGTGAAAGAAAAGGTAAGCTGATCAGA | |
| 19026 reads | ATTTTTCTTTTTTCTAGGGC------ | ... | ----------------GAAAAGGTAAGCTGATCAGA | 0 |
| 208 reads | ATTTTTCTTTTTTCTAGGGC------ | ... | ----------------GAAAAGGTAAGCTGATCAGA | 1 |
| 112 reads | ATTTTTCTTTTTTCTAGGGC------ | ... | ----------------GAAAAGGTAAGCTGATCAGA | 1 |
| | | (79 bases omitted) | | |

Reverse sequencing results

| | | | | |
|---|---|---|---|---|
| Reference Sequence | ATTTTTCTTTTTTCTAGGGCAAGAGG | ... | ATTTGAAAACGTGAAAGAAAAGGTAAGCTGATCAGA | |
| 17915 reads | ATTTTTCTTTTTTCTAGGGC------ | ... | ----------------GAAAAGGTAAGCTGATCAGA | 0 |
| 88 reads | ATTTTTCTTTTTTCTAGGGC------ | ... | ----------------GAAAAGGTAAGCTGATCAGA | 1 |
| 87 reads | ATTTTTCTTTTTTCTAGGGC------ | ... | ----------------GAAAAGGTAAGCTGATCAGA | 2 |
| | | (79 bases omitted) | | |

**b**

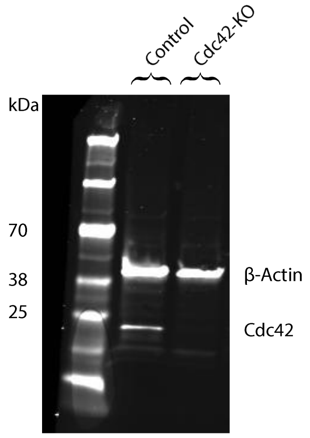


**Supplementary Figure 3:** The Cdc42 knockout is homozygous. (**a**) Amplicon sequencing results. The dominant sequence was identical for both forward and reverse primers. The single dominant read indicates that the same deletion is present for both alleles. (**b**) Full-lane western blot. Top bands correspond to an anti-β-Actin antibody while bottom band is reactive to an anti-Cdc42 antibody.

Supplementary Figure 4

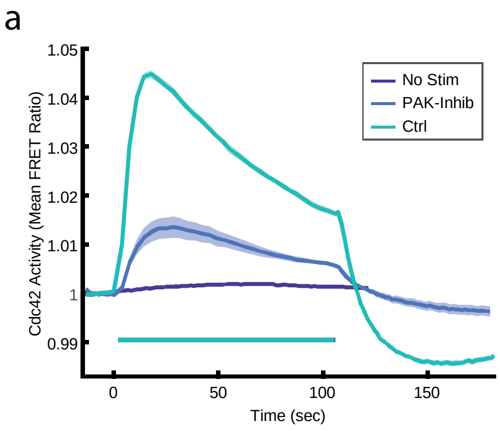


**Supplementary Figure 4:** Inhibition of PAK1 reduces Cdc42 response magnitude regardless of stimulation light power. (**a**) Control and PAK1-Inhibited cells responding to medium power light stimulation (Relative light intensity = 50). Shaded error region ± s.e.m. of $n_{\text{well replicates}}$ = 65 for non-stimulated, $n_{\text{well replicates}}$ = 59 for control and $n_{\text{well replicates}}$ = 16 for PAK1 inhibited cells.

Supplementary Figure 5

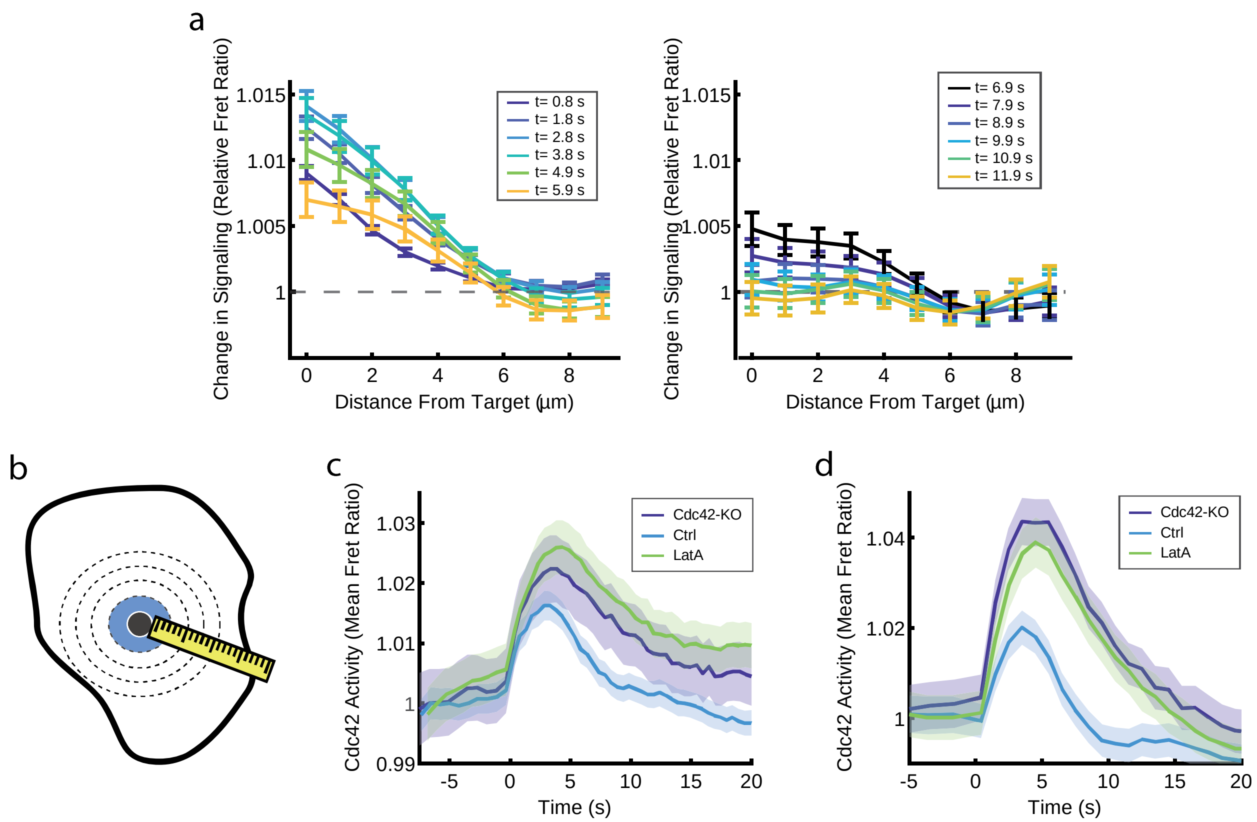


**Supplementary Figure 5:** The Cdc42-KO and Latrunculin-A perturbations prolong the duration of the Cdc42 response. (**a**) Time course encompassing activation and attenuation of Cdc42 response to single pulse center stimulation experiment. Relative Cdc42 response as a function of distance from the stimulation target site for control cells. The response attenuates by ~10 seconds post stimulation (Right panel). Experimental stimulation condition: one, 0.8 µW light-pulse with 10ms duration. Error bars represent ± s.e.m. of n=181 cells. (**b**) Center stimulation experiment schematic. Pixels < 1µm from the stimulation site (blue shaded region) were used to generate plots in (**c-d**). (**c**) Cdc42 activity as a function of time for control, Cdc42-KO, and Latrunculin-A treated cells responding to one, 0.8 µW light-pulse with 10 ms duration. Error bars represent ± s.e.m. of n=181 cells for control, n=67 for Cdc42-KO, and n=175 for Latrunculin-A. (**d**) Cdc42 activity as a function of time for control, Cdc42-KO, and Latrunculin-A treated cells responding to one, 4.3 μW light-pulse with 10 ms duration. Error bars represent ± s.e.m. of n=131 cells for control, n=43 for Cdc42-KO, and n=105 for Latrunculin-A-treated cells.

Supplementary Figure 6

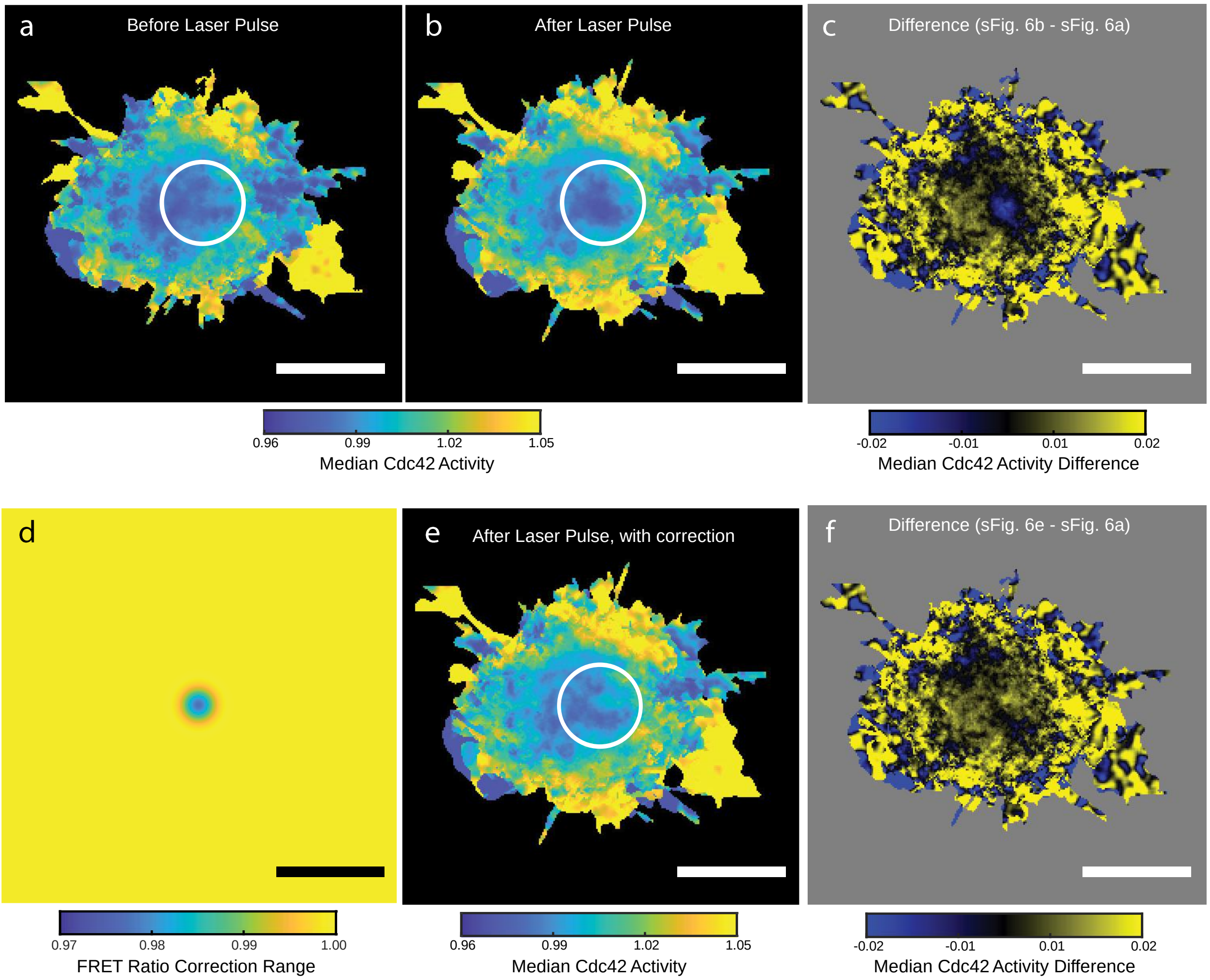


**Supplementary Figure 6:** Photobleaching correction for bleaching due to FRAP laser stimulation. (**a-b**) Median Cdc42 FRET ratio of n=79 cells for the image immediately preceding (**a**) and succeeding (**b**) stimulation. Cells were stimulated with a strong, 37 µW, 10 ms laser pulse to maximize bleaching for computing the correction. White circles indicate the FRAP target region. Photobleaching causes a reduction in FRET signal (b). (**c**) Difference between post-stimulus and pre-stimulus image. (**d**) Photobleaching correction image for the frame immediately succeeding FRAP stimulation. (**e**) Median Cdc42 FRET ratio of n=79 cells where the photobleaching correction was applied to the post-stimulation frame. (**f**) Difference between photobleaching corrected post-stimulus and pre-stimulus images. Scale bar for all panels, 15 µm.

**Supplementary Video 1:** Global stimulation of Parapinopsina induced Cdc42 activity is dependent on UV light stimulation and the presence of the 9-*cis*-retinal cofactor (as in Fig. 1d). Scale bar, 75 µm. Raw fluorescence intensity is shown on the left, and FRET ratio is shown on the right. Representative movies for unstimulated cells, cell stimulated without the retinal cofactor, and for stimulation of cells with the full optogenetic system were appended.

**Supplementary Video 2:** Localized optogenetic stimulation can drive a chemotaxis-like response (as in Fig. 1e). Cdc42 activity is localized to the leading edge in optogenetically driven cells, matching the spatial activity pattern seen in chemotaxing neutrophils. Scale bar, 25 µm. Color bar for both videos represents Cdc42 activity. Raw fluorescence intensity is shown on the left, and FRET ratio is shown on the right. Movies for two representative cells were appended.

**Supplementary Video 3:** Cdc42 activity is dose-dependent on receptor input strength (as in Fig. 2b). Scale bar, 75 µm. Representative movies for populations of cells stimulated with different light intensities were appended.

**Supplementary Video 4:** Cdc42-KO cells engage in many repolarization events, including building large, unstable lamellipodia. Scale bar, 15 µm. Movies of a representative control cell and Cdc42-KO cells were appended.

**Supplementary Video 5:** Cdc42-KO cells demonstrating the cytoplasmic tether phenotype; the cell front pulls away from the cell body, but remains attached via a thin cytoplasmic tether. Scale bar, 15 µm.

**Supplementary Video 6:** Cdc42 spatial response to local, stimulation in the cell center (as in Fig. 5a). A single, 4.3 μW, 10 ms laser pulse was applied at t=0. Purple circle indicates the stimulation site. Color bar represents Cdc42 activity. Scale bar, 25 μm.
